# Degradable porous PLGA/PCL membrane enable a lung alveoli-on-a-chip for modeling particulate-induced alveolar injury

**DOI:** 10.64898/2026.04.03.716399

**Authors:** Jae-Won Choi, Sikandar Azam, MyAnh Hisaeda, Shimin Liu, Si-Yang Zheng

## Abstract

Understanding how airborne particulates disrupt the alveolar barrier requires *in vitro* systems that recapitulate both the structure and transport properties of the lung air-blood interface. Here, we report a biodegradable lung alveoli-on-a-chip enabled by porous poly(lactic-co-glycolic acid)/polycaprolactone (PLGA/PCL) membranes with an interconnected porous architecture generated via porogen-assisted phase separation process. The membrane exhibits tunable degradation behavior, allowing progressive increases in surface porosity (∼40%) and reduction in thickness (∼3 µm) during culture, while PCL maintains mechanical integrity under dynamic conditions. These degradation-driven structural changes regulate membrane transport properties, leading to enhanced permeability and supporting the formation of a functional epithelial-endothelial barrier under air-liquid interface (ALI) culture with breathing-mimetic cycling strain. Primary human alveolar epithelial and microvascular endothelial cells formed confluent, junctional monolayers on opposing membrane surfaces, exhibiting stable barrier function and high viability throughout the culture period. As a functional application, the platform was used to assess diesel particulate matter (DPM)-induced alveolar injury. Apical exposure to DPM induced dose-dependent cytotoxicity, increased barrier permeability, elevated reactive oxygen species, and DNA damage in both epithelial and endothelial layers, demonstrating trans-barrier propagation of particulate-induced injury. Pharmacological modulation with roflumilast-N-oxide (RNO), a phosphodiesterase-4 (PDE4) inhibitor, selectively attenuated oxidative stress and inflammatory responses, with limited effects on barrier integrity. Together, this work establishes degradable PLGA/PCL membranes as tunable interface materials for lung-on-a-chip systems, where structural evolution during degradation directly governs transport and barrier function. The resulting platform provides a physiologically relevant approach for studying particulate toxicity and therapeutic modulation at the alveolar interface.

## 1. Introduction

Exposure to ambient particulate matter (PM) is a major contributor to the global burden of disease, with extensive epidemiological evidence linking it to increased morbidity and mortality from cardiovascular and respiratory disorders [1, 2]. Among anthropogenic PM sources, diesel particulate matter (DPM) is of particular concern due to its widespread emission from transportation and industrial activities and its propensity for deep lung deposition [3–5]. DPM consists of a carbonaceous core enriched with transition metals and particle-bound organic compounds, including polycyclic aromatic hydrocarbons (PAHs), which are known to induce oxidative stress, inflammation, and genotoxic damage in pulmonary tissues [3]. As a result, diesel exhaust has been classified as a Group 1 carcinogen by the International Agency for Research on Cancer (IARC) [5]. Beyond pulmonary toxicity, inhalation of DPM has also been strongly associated with systemic cardiovascular complications, including endothelial dysfunction, atherosclerosis, and increased risk of myocardial infarction, highlighting the ability of inhaled particles to perturb both local alveolar barriers and distal vascular systems. These broad and multi-organ health impacts underscore the critical need for human-relevant models that can resolve how DPM disrupts the alveolar epithelial–endothelial interface, where inhalation exposure is first translated into both pulmonary and systemic pathophysiology [6–8]. Although epidemiological studies have firmly established the association between DPM exposure and adverse respiratory and systemic outcomes [9–11], the cellular and barrier-level mechanisms underlying DPM-induced alveolar injury are not yet fully elucidated.

A major barrier to mechanistic insight is the lack of physiologically relevant *in vitro* models that faithfully recapitulate the structural and functional features of the human alveolar interface. Conventional cell culture systems, including standard monocultures and Transwell® assays, fail to reproduce key features of the alveolus, such as the ultrathin epithelial-endothelial barrier, air-liquid interface (ALI) exposure, and breathing-associated mechanical deformation [12, 13]. While animal models provide valuable insights into pulmonary toxicity, species-specific differences and limited accessibility to layer-resolved cellular responses constrain their translational relevance [14]. These limitations highlight the need for human-relevant experimental platforms capable of dissecting particulate-induced injury at the alveolar-capillary interface.

Organ-on-a-chip technologies have emerged as powerful tools for reconstructing tissue microenvironments by integrating microfluidics, biomaterials, and dynamic mechanical cues [15, 16]. In particular, lung-on-a-chip systems have enabled the study of airway and alveolar physiology under more realistic conditions [17–19]. However, a central limitation of many existing platforms lies in the membrane separating epithelial and endothelial compartments. Commonly used synthetic membranes are non-degradable, relatively thick, and exhibit limited pore interconnectivity, restricting transport and failing to capture the dynamic structural remodeling of the native alveolar interstitium [17–22]. These constraints can alter barrier function and diminish physiologically relevant epithelial-endothelial crosstalk.

To address these limitations, we developed a biodegradable lung alveoli-on-a-chip incorporating a porous poly(lactic-co-glycolic acid)/polycaprolactone (PLGA/PCL) membrane as a tunable interface scaffold. The membrane exhibits an interconnected porous architecture and undergoes progressive degradation during culture, resulting in increased membrane porosity and reduced thickness over time. This degradation-driven structural evolution enhances transport across the interface while the PCL component preserves mechanical integrity, allowing sustained breathing-mimetic cyclic deformation under ALI conditions.

Using this platform, we systematically investigated the effects of DPM exposure at the epithelial-endothelial interface. Apical exposure to DPM induced dose-dependent cytotoxicity, barrier disruption, oxidative stress, and DNA damage in both epithelial and endothelial layers, revealing trans-barrier propagation of particulate-induced injury. Beyond toxicological assessment, the ability to evaluate therapeutic modulation remains an important need in *in vitro* lung models. Phosphodiesterase-4 (PDE4) inhibitors, such as roflumilast, are clinically approved anti-inflammatory agents used in the treatment of chronic obstructive pulmonary disease (COPD) [23, 24]. Its active form, roflumilast N-oxide (RNO), exerts anti-inflammatory effects through cAMP-mediated signaling pathways [25]. Here, we further demonstrate the utility of this platform for evaluating pharmacological intervention against particulate-induced injury. Administration of RNO to the vascular compartment selectively attenuated DPM-induced oxidative stress and inflammatory cytokine production, while exerting minimal effects on cell viability and barrier permeability.

Together, this work establishes degradable porous membranes as functional interfacial materials for lung-on-chip systems, where structural evolution during degradation modulates transport and barrier behavior. The resulting platform provides a physiologically relevant approach for dissecting particulate-toxicity and evaluating therapeutic modulation at the alveolar interface.

## 2. Results

### 2.1 Conceptual overview and fabrication of porous PLGA/PCL membranes for lung alveoli-on-a-chip

We developed a lung alveoli-on-a-chip platform incorporating a biodegradable porous membrane to recapitulate key structural and functional features of the human alveolar interface. *In vivo*, the air-blood barrier consists of an alveolar epithelial layer and a capillary endothelial layer separated by an ultrathin interstitial region (∼1–2 µm), while undergoing cyclic deformation during respiration (figure 1(A)) [26]. To mimic this physiological architecture, we engineered a biodegradable, mechanically compliant membrane using two polymers with complementary properties. Poly(lactic-co-glycolic acid) (PLGA) was selected for its rapid hydrolytic degradation, which enables time-dependent increases in membrane porosity and reductions in thickness [27]. Polycaprolactone (PCL), in contrast, exhibit slower degradation and enhanced elasticity, thereby providing mechanical stability and allowing repeated deformation under breathing-like loading [28]. By combining these materials, the membrane was designed to exhibit both structural evolution and mechanical resilience during culture.

**Figure 1.**
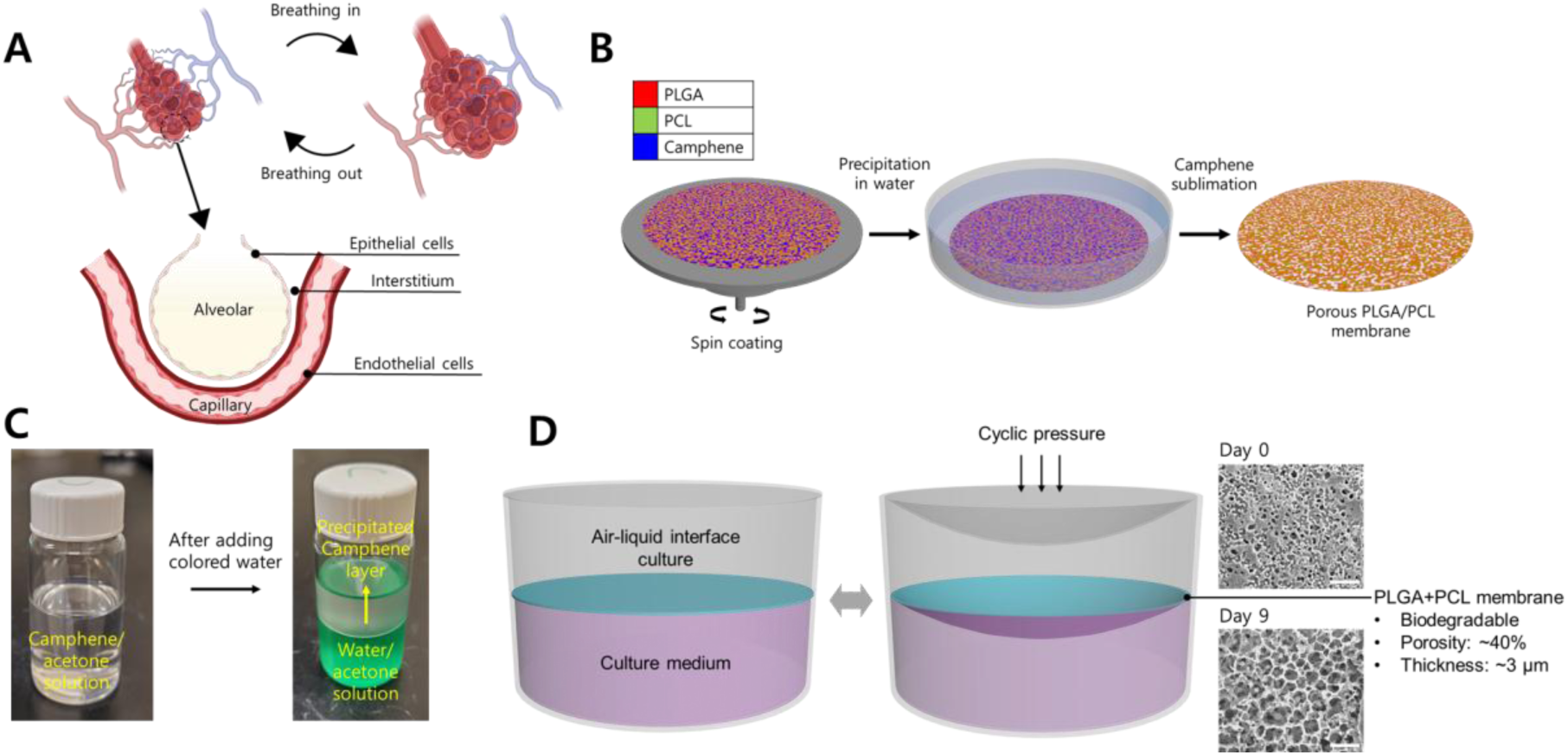
Design and fabrication of porous PLGA/PCL membranes for lung alveoli-on-a-chip. (**A**) Schematic of the alveolar-capillary interface, consisting of an epithelial layer and an endothelial layer separated by an ultrathin interstitial region (∼1–2 µm) that undergoes cyclic deformation during breathing. (**B**) Schematic illustration of the fabrication process of porous PLGA/PCL membranes via porogen-assisted nonsolvent-induced phase separation. (**C**) Optical images of the camphene/acetone solution (left) and the formation of precipitated camphene layer upon mixing with colored water (right), illustrating the phase separation and porogen templating process. (**D**) Schematic of the lung alveoli-on-a-chip system under air-liquid interface (ALI) cultures with breathing-mimetic cyclic deformation. Insets are representative SEM images of the membrane at Day 0 and Day 9, showing increased porosity during degradation. Scale bars: 20 µm.

The porous architecture was generated using a porogen-assisted nonsolvent-induced phase separation (NIPS) process (figure 1(B)) [29]. Briefly, a homogeneous PLGA/PCL-camphene solution in acetone was spin-coated onto the substrate and subsequently immersed in water, which acts as a nonsolvent for both polymers and camphene. Rapid solvent exchange drove polymer phase separation and solidification, while camphene crystallized within the polymer matrix to form a dendritic framework (figure 1(C)) [30]. Following removal of camphene by sublimation produced an interconnected porous structure throughout the membrane.

The resulting PLGA/PCL membrane was integrated between the apical and basolateral chambers of a PDMS-based device, defining the epithelial-endothelial interface (figure 1(D)). Epithelial cells were cultured on the apical surface under air exposure, while endothelial cells were cultured on the basolateral side under medium, enabling air-liquid interface (ALI) co-culture. During culture, the membrane progressively degraded, resulting in an increase in surface porosity to over 40% and a reduction in thickness to approximately 3 µm over 9 days. At the same time, the PCL component preserved sufficient mechanical integrity to support cyclic deformation. Breathing-mimetic mechanical stimulation (10% linear strain) was applied to reproduce physiological strain conditions.

### 2.2 Optimization and characterization of porous PLGA/PCL membranes with different blend ratios

To identify an optimal membrane composition that balances porosity evolution, mechanical compliance, and degradation stability, porous PLGA/PCL membranes with different blend ratios (PLGA:PCL = 10:0, 9:1, 8:2, and 7:3) were systematically characterized. Before degradation, the surface porosity of the membranes was comparable across all PLGA/ PCL compositions, showing no substantial dependence on blend ratio (10:0, 23.4%; 9:1, 20.1%; 8:2, 22.6%; and 7:3, 23.7%) (figure 2(A) and (B)). Upon incubation in phosphate-buffered saline (PBS) at 37 °C, the membrane underwent composition-dependent structural evolution. After 9 days, surface porosity increased significantly for all formulations, with a more pronounced increase observed in PLGA-rich membranes (10:0, 48.8%; 9:1, 46.7%; 8:2, 40.1%; and 7:3, 37.1%). This trend is consistent with the much faster hydrolytic degradation of PLGA relative to PCL, leading to enhanced pore enlargement in membranes with higher PLGA content. Concurrently, membrane thickness decreased significantly across all compositions, from 5.75 to 3.14 µm for 10:0, from 5.8-6.1 µm to 3.2-3.7 µm after degradation (figure 2(C)).

**Figure 2.**
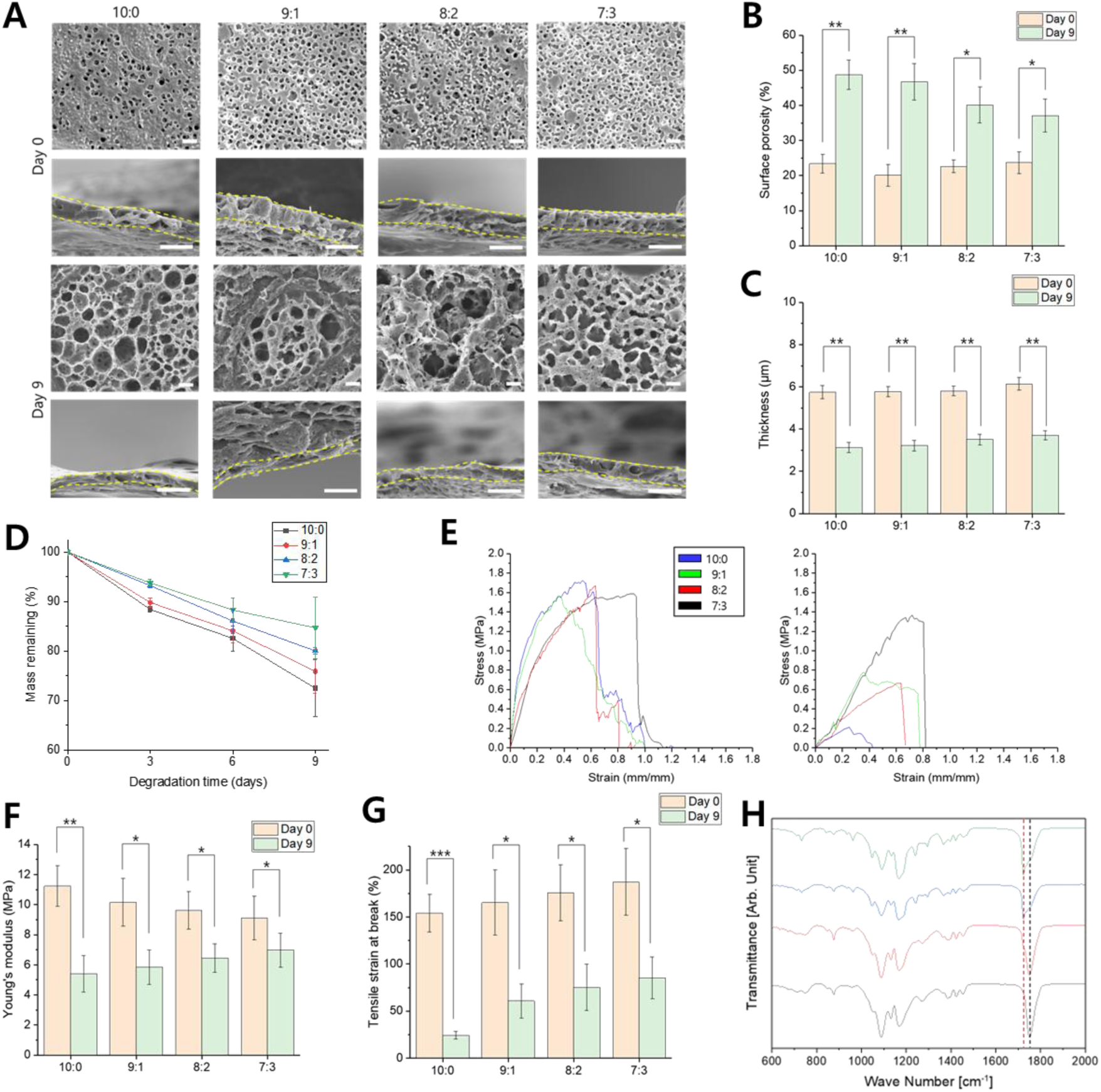
Composition-dependent structural, degradation, and mechanical properties of porous PLGA/PCL membranes. **(A**) Top-view and cross-sectional SEM images of membranes with different PLGA:PCL ratios (10:0, 9:1, 8:2, and 7:3) before and after degradation in PBS (37 °C) for 9 days. Scale bars: 10 µm. (**B**) Mass loss of porous PLGA/PCL membranes with different blend ratios during degradation (n = 4). (**C**) Surface porosity of membranes with before and after degradation (n = 4). (**D**) Membrane thickness before and after degradation (n = 4). (**E**) Representative stress-strain curves of membranes before (left) and after degradation (right) (n = 4). (**F**) Young’s modulus of membranes before and after degradation (n = 4). (**G**) Tensile strain at break of membranes before and after degradation (n = 4). (**H**) FT-IR spectra confirming PLGA/PCL composition (10:0, black; 9:1, red; 8:2, blue; and 7:3, green), showing characteristic carbonyl peaks for PLGA (1750 cm⁻¹) and PCL (1722 cm⁻¹).

Mass-loss measurements provided additional quantitative evidence for the progressive degradation behavior of the composite membrane. Pure PLGA membranes showed substantial mass loss (27.5% after 9 days), while incorporation of PCL reduced the overall degradation rate, with the 7:3 composition showing markedly lower mass loss (15.3%) (figure 2(D)). These results demonstrate that PCL effectively moderates degradation kinetics while maintaining the porous architecture.

Mechanical properties were evaluated to assess membrane integrity before and after degradation (figure 2(E)). Prior to degradation, increasing PCL content reduced the Young’s modulus while increasing ductility (strain at break), reflecting the greater elasticity of PCL (figure 2(F) and (G)). After 9 days of degradation, PLGA underwent substantial degradation, whereas PCL remained largely stable, preserving mechanical strength. As a result, membranes with higher PCL content showed attenuated reductions in both Young’s modulus and strain at break, indicating improved mechanical stability during degradation.

Fourier-transform infrared (FT-IR) spectroscopy confirmed the successful incorporation of both polymers in the composite membrane. With increasing PCL content, the characteristic carbonyl stretching peak at 1722 cm⁻¹ became more prominent, whereas pure PLGA membrane exhibited a single peak at 1750 cm⁻¹ without detectable PCL-associated features (figure 2(H)).

Based on these results, a PLGA:PCL blend ratio of 8:2 was selected for subsequent experiments, as it provides a balanced combination of degradation-driven porosity enhancement and mechanical stability, enabling both effective transport and structural integrity during culture.

### 2.3 Biocompatibility and barrier integrity of PLGA/PCL membrane-based lung alveoli-on-a-chip

To support epithelial cells under ALI conditions, the membrane must permit transport of aqueous media from the basolateral side to sustain nutrient supply. As shown in figure 3(A), a water-soluble dye readily permeated through the PLGA/PCL membrane, confirming its permeability to aqueous transport. The pristine membrane displayed a water contact angle of 90.8°, indicating moderate hydrophobicity (figure 3(B)). Following gelatin coating, the contact angle decreased to 62.2°, reflecting enhanced surface wettability favorable for cell attachment.

**Figure 3.**
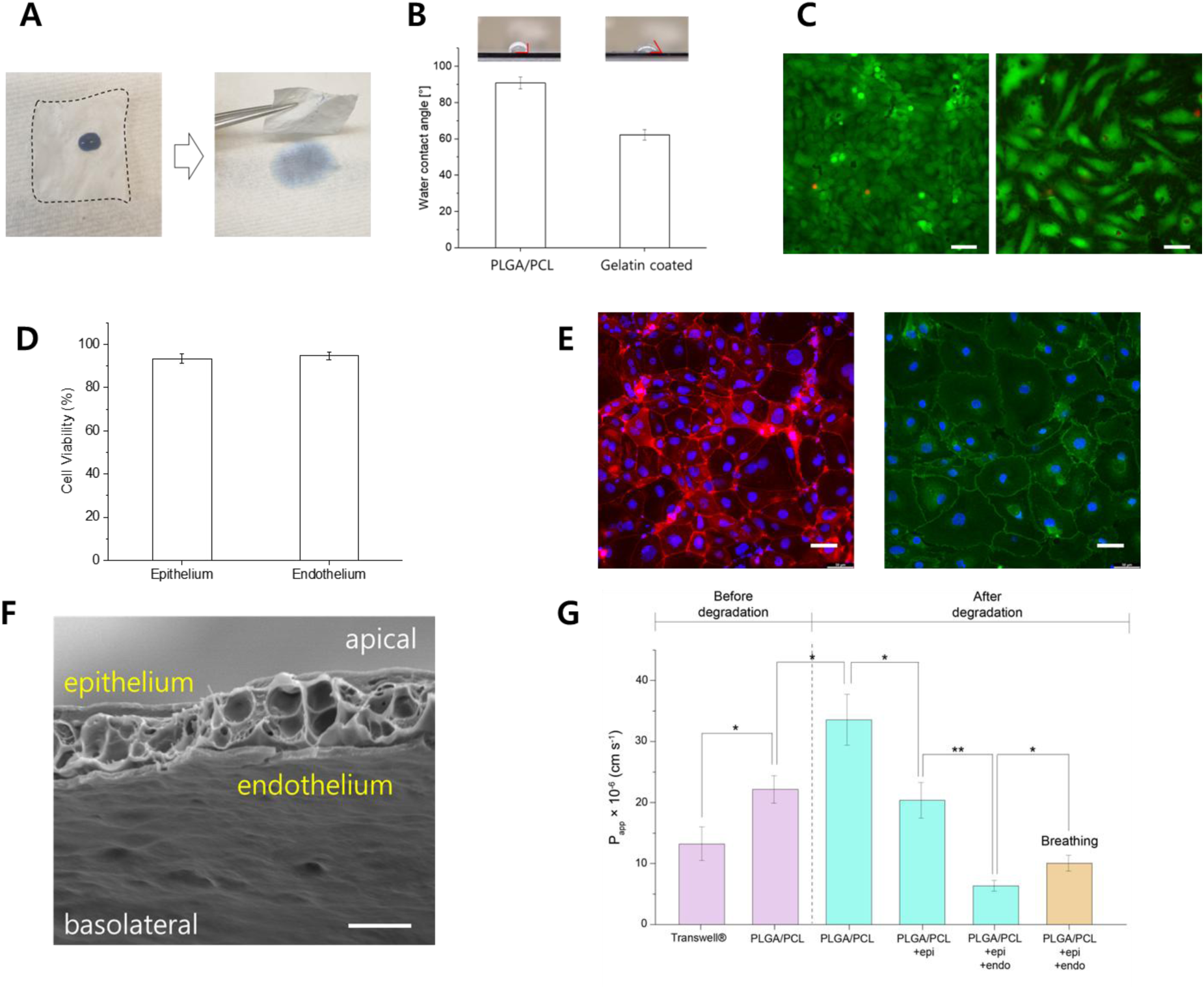
Biocompatibility and barrier function of PLGA/PCL membrane-based lung alveoli-on-a-chip. (**A**) Optical images of a water-soluble dye droplet on the membrane before and after permeation, demonstrating aqueous transport through the porous structure. (**B**) Water contact angle measurements of membranes before and after gelatin coating (n = 4). (**C**) Representative Live/Dead fluorescence images of HPAECs and HMVECs co-cultured on the membrane. (**D**) Quantification of cell viability for epithelial and endothelial co-culture (n = 4). (**E**) Immunofluorescence images showing tight junction formation in HPAECs (ZO-1, red) and adherens junction formation in HMVECs (VE-cadherin, green). Scale bars: 50 µm. (**F**) Cross-sectional SEM image of epithelial-endothelial co-culture on the porous membrane under ALI conditions. Scale bar: 3 µm. (**G**) Apparent permeability (P_app_) of 4 kDa dextran across Transwell® and PLGA/PCL membranes before (purple) and after degradation (teal), and across epithelial and epithelial-endothelial barriers formed on degraded membranes under static and cyclic stretching conditions (orange) (n = 4).

Epithelial-endothelial co-culture was established by first seeding human lung microvascular endothelial cells (HMVECs) onto the basolateral surface of the membrane. After 3 hours of incubation to allow cell attachment, the device was inverted and human primary alveolar epithelial cells (HPAECs) were seeded on the apical surface. Co-culture were maintained under liquid-liquid culture conditions for 7 days to promote barrier formation, after which the apical medium was removed to initiate ALI culture. Breathing-mimetic cyclic stretching was applied 24 hours after the transition to ALI.

After 9 days of culture, Live/Dead staining demonstrated high cell viability for both HPAECs and HMVECs (figure 3(C)), with quantitative analysis showing excellent cell viability of 93.4% and 94.7% (figure 3(D)), respectively. Under ALI conditions with cyclic stretching, both cell types formed confluent monolayers, as evidenced by ZO-1-positive tight junctions in epithelial cells and VE-cadherin-positive adherens junctions in endothelial cells (figure 3(E)). Cross-sectional SEM imaging further revealed distinct epithelial and endothelial layers on opposite sides of the ∼3 µm-thick membrane, demonstrating stable co-culture and structural integrity under dynamic mechanical loading (figure 3(F)).

The transport properties of the membrane were evaluated by measuring apparent permeability (P_app_). Compared with 0.4 µm pore-sized Transwell® membranes, the PLGA/PCL membrane exhibited a 1.68-fold higher permeability, indicating enhanced transport capacity and improved potential for epithelial-endothelial crosstalk (figure 3(G)) [31, 32]. Following degradation, permeability increased by 1.50-fold, consistent with the observed increase in porosity. Upon formation of cellular barriers on the degraded membrane, permeability decreased to 20.35 × 10⁻^6^ cm·s⁻¹ for epithelial monoculture and 6.41 × 10⁻^6^ cm·s⁻¹ for epithelial-endothelial co-culture, confirming the formation of functional barrier properties [33, 34]. Under these conditions, cyclic stretching increased permeability by 1.57-fold, reflecting dynamic modulation of barrier transport by breathing-associated mechanical strain.

### 2.4 Toxicity assessment of diesel particulate matter in a lung alveoli-on-a-chip

To evaluate the functional capability of the platform for particulate toxicity assessment, diesel particulate matter (DPM) was applied to the apical surface of the lung alveoli-on-a-chip under ALI conditions (figure 4(A)). The DPM particles exhibited a broad size distribution (0.1–10 μm), representative of inhalable particulates (figure 4(B)). The apparent discrepancy between Zetasizer (dynamic light scattering, DLS) and Mastersizer (laser diffraction) measurements reflects their distinct measurement principles: DLS reports hydrodynamic diameters weighted toward smaller particles in suspension, while laser diffraction provides volume-weighted distributions that emphasize larger particles. Together, these complementary techniques capture the polydispersity nature of DPM. To induce measurable biological responses within a practical experimental timeframe, DPM was administered at surface doses ranging from 0 to 60 µg/cm².

**Figure 4.**
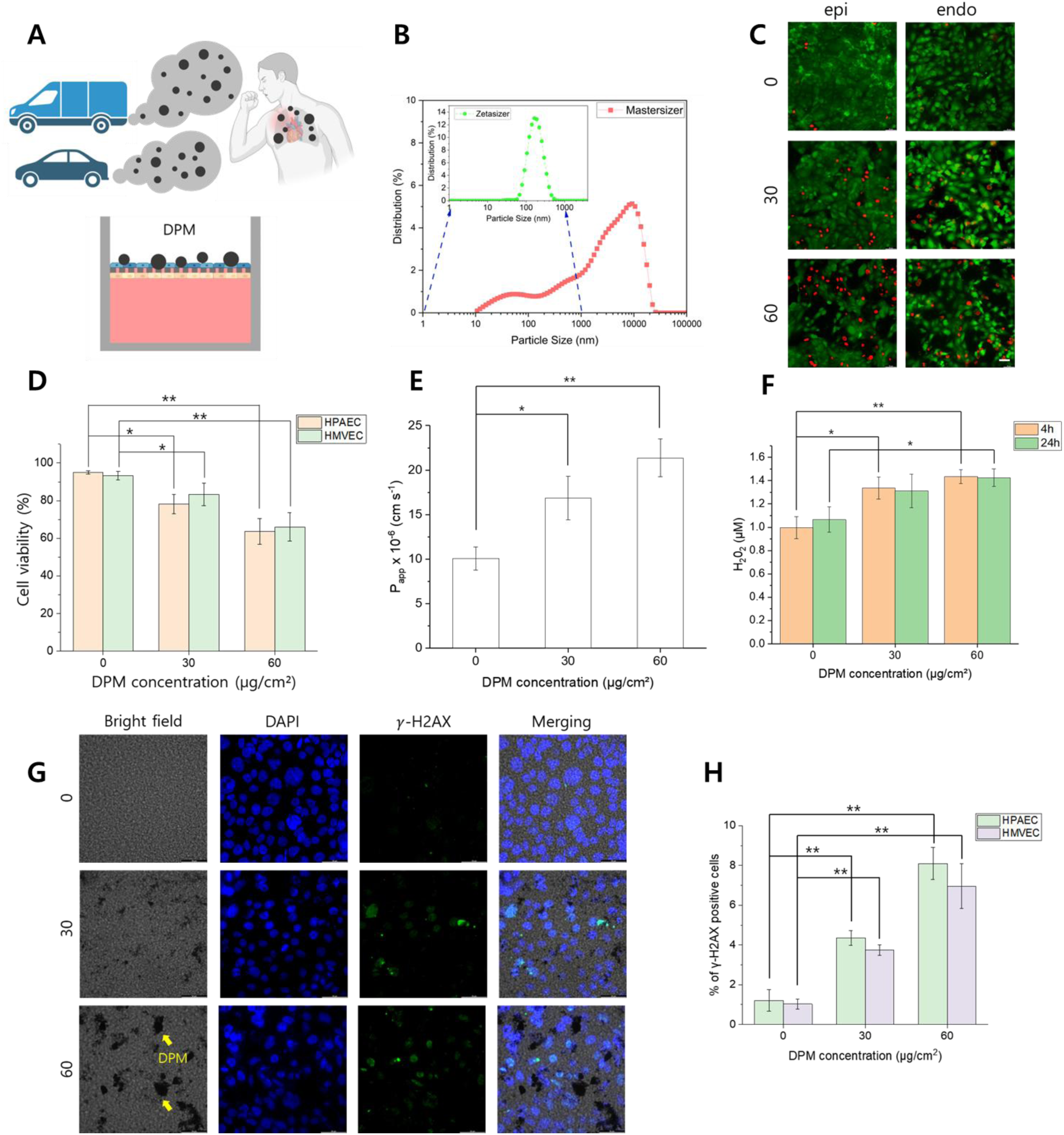
Assessment of DPM-induced toxicity in a lung alveoli-on-a-chip. (**A**) Schematic of the lung alveoli-on-a-chip for apical exposure to DPM under ALI conditions. (**B**) Particle size distribution of DPM measured by dynamic light scattering (Zetasizer) and laser diffraction (Mastersizer), highlighting the polydisperse nature of the particles. (**C**) Live/Dead fluorescence images of epithelial (HPAEC) and endothelial (HMVEC) cells after 24-hour exposure to DPM at 0, 30, and 60 µg/cm². Scale bar: 50 µm. (**D**) Quantification of cell viability in epithelial and endothelial layers following DMP exposure. (**E**) Apparent permeability (P_app_) of the 4 kDa FITC-dextran across the epithelial-endothelial barrier measured at 24 hours after DPM exposure (n = 4). (**F**) Reactive oxygen species (ROS) levels quantified by hydrogen peroxide (H₂O₂) produced after 24 hours after exposure (n = 4). (**G**) Immunofluorescence images of γ-H2AX staining showing DNA double-strand breaks in epithelial cells after DPM exposure. Scale bar: 50 µm. (**H**) Quantification of γ-H2AX-positive epithelial and endothelial cells following DPM exposure (n = 4).

DPM exposure resulted in a dose-dependent reduction in cell viability in both epithelial and endothelial layers (figure 4(C) and (D)). At 30 µg/cm², epithelial and endothelial viabilities decreased to 78.2% and 83.3%, respectively, and further declined to 63.6% and 66.0% at 60 µg/cm². The comparable reduction in endothelial viability indicates that DPM particles traverse the epithelial layer and porous membrane, thereby exerting downstream effects on the vascular compartment. This interpretation is further supported by quantitative measurements showing that a substantial fraction of DPM translocates across the membrane within 24 hours (figure S1).

Barrier function was assessed by measuring apparent permeability. As the DPM concentration increased, permeability was progressively elevated, indicating disruption of epithelial-endothelial integrity (figure 4(E)). In parallel, oxidative stress was evaluated by quantifying hydrogen peroxide-based reactive oxygen species (ROS), which showed a concentration-dependent increase following DPM exposure (figure 4(F)).

Genotoxic effects were assessed by immunostaining for the phosphorylated form of histone H2AX (γ-H2AX), a marker of DNA double-strand breaks. As shown in figure 4(G), the fraction of γ-H2AX-positive cells increased with DPM concentrations in both epithelial and endothelial layers. Baseline levels of DNA damage were minimal (HPAEC: 1.2%; HMVEC: 1.0%), whereas exposure to 60 µg/cm² DPM increased γ-H2AX positivity to 8.1% in epithelial cells and 7.0% in endothelial cells (figure 4(H)).

### 2.5 Mitigation of DPM-induced alveolar toxicity by roflumilast-N-oxide (RNO) in a lung alveoli-on-a-chip

Roflumilast is a selective phosphodiesterase-4 (PDE4) inhibitor that is metabolized *in vivo* into its active form, roflumilast N-oxide (RNO) [23], which elevates intracellular cyclic AMP (cAMP) levels and modulates inflammatory signaling pathways [23–25]. To evaluate the potential of the platform for therapeutic assessment, RNO (100 nM) was administered to the basolateral chamber following apical exposure to DPM.

Live/Dead imaging showed that DPM exposure induced cell death in both epithelial and endothelial layers, with no apparent rescue upon RNO co-treatment (figure 5(A)). Quantitative analysis confirmed that RNO did not significantly improve overall cell viability (figure 5(B)). Similarly, measurement of permeability across the epithelial-endothelial barrier revealed that RNO exerted only a marginal effect on DPM-induced barrier disruption, with no statistically significant restoration of barrier integrity (figure 5(C)).

**Figure 5.**
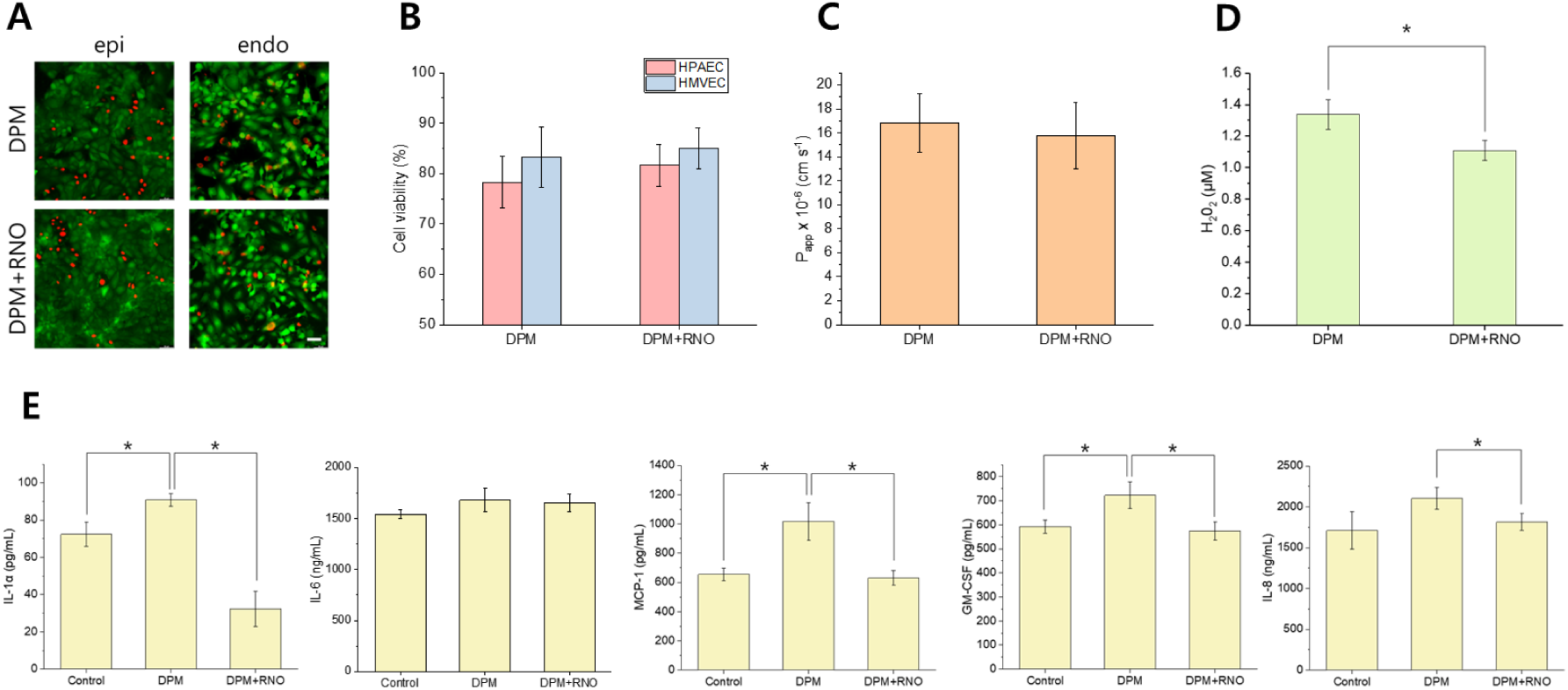
Mitigation of DPM-induced alveolar toxicity by roflumilast-N-oxide (RNO) in a lung alveoli-on-a-chip. (**A**) Representative Live/Dead fluorescence images of epithelial and endothelial cells following exposure to DPM (30 µg/cm²) with or without RNO treatment (100 nM). Scale bar: 50 µm. (**B**) Quantification of cell viability in epithelial and endothelial layer under indicated conditions. (**C**) Apparent permeability (P_app_) of the epithelial-endothelial barrier following DPM exposure in the absence or presence of RNO (n = 4). (**D**) ROS levels quantified by H₂O₂ production after DPM exposure with or without RNO treatment. (**E**) Cytokine secretion profiles measured from basolateral media 24 hours after DPM exposure in the absence or presence of RNO (n = 3).

In contrast, RNO markedly modulated oxidative and inflammatory responses. Hydrogen peroxide measurements demonstrated a significant reduction in intracellular ROS levels in RNO-treated chips compared with DPM-only conditions (figure 5(D)). Furthermore, multiplex cytokine analysis of basolateral media revealed that RNO attenuated the secretion of multiple pro-inflammatory cytokines induced by DPM exposure (figure 5(E)). These observations are consistent with the known anti-inflammatory effects of PDE4 inhibition in pulmonary cells [35–39].

Collectively, these findings suggest that RNO selectively mitigates DPM-induced oxidative stress and inflammatory signaling, while exerting limited effects on cell viability and barrier function within the experimental timeframe. This differential response highlights the ability of the platform to resolve distinct aspects of particulate-induced injury and therapeutic modulation at the alveolar interface.

## 3. Discussion

This study presents a biodegradable membrane-enabled lung alveoli-on-a-chip that advances *in vitro* modeling of particulate-induced lung injury by integrating dynamic structural evolution, physiologically relevant transport, epithelial-endothelial co-culture under ALI conditions, and breathing-mimetic cyclic deformation. A central innovation of this work lies in the use of a degradable PLGA/PCL membrane as a tunable interface material. Unlike conventional non-degradable membranes, the PLGA component undergoes controlled hydrolytic degradability, leading to progressively increases in porosity and reduction in thickness, while the PCL component preserves mechanical integrity under cyclic deformation. This combination enables a time-evolving interface whose structural and transport properties more closely approximates those of the native alveolar barrier.

The degradation-driven evolution of membrane structure has direct functional consequences. Increased porosity and reduced thickness enhance permeability and facilitate epithelial-endothelial crosstalk, while maintaining sufficient mechanical stability to support breathing-mimetic cyclic strain. As a result, the platform supports the formation of a functional alveolar barrier with physiologically relevant transport characteristics. This dynamic structure-function relationship distinguishes the present system from static membrane-based models, in which fixed pore architecture and thickness limit transport and mechanotransduction.

According to global disease burden assessments, particulate matter is recognized as a major environmental risk factor for cardiovascular and respiratory diseases and is ranked as the second leading global risk factor for premature mortality and disability. Among anthropogenic PM sources, DPM is of particular concern due to its widespread emission from transportation and its high likelihood of deep lung deposition. DPM consists of a carbonaceous core enriched with transition metals and particle-bound organic compounds such as polycyclic aromatic hydrocarbons (PAHs), which are known to induce oxidative stress, inflammation, and genotoxic damage in pulmonary tissues upon inhalation, and has therefore been classified as a Group 1 carcinogen by the International Agency for Research on Cancer (IARC). Despite strong epidemiological links between DPM and respiratory disease, the lack of physiologically relevant *in vitro* models that recapitulate the alveolar microenvironment has hindered mechanistic, layer-resolved understanding of DPM-induced alveolar injury.

Using this lung alveoli-on-a-chip platform, we demonstrate that DPM induces dose-dependent alveolar injury characterized by cytotoxicity, barrier disruption, oxidative stress, and DNA damage in both epithelial and endothelial layers. Notably, comparable reductions in epithelial and endothelial viability indicates trans-barrier propagation of particulate-induced injury, highlighting the importance of modeling the coupled epithelial-endothelial interface. These findings underscore the value of incorporating both high permeability and dynamic mechanical cues to capture physiologically relevant responses that are not readily observed in conventional *in vitro* systems.

The platform also enables the evaluation of pharmacological modulation in a layer-resolved manner. Treatment with RNO revealed a selective protective profile, with significant attenuation of oxidative stress and pro-inflammatory cytokine production but minimal effects on cell viability and barrier integrity. This differential response indicates that DPM-induced injury comprises multiple components that are pharmacologically separable, where biochemical signal pathways associated with oxidative damage and inflammation can be effectively modulated without immediate restoration of structural barrier function. These findings suggest that anti-inflammatory interventions alone may be insufficient to reverse particulate-induced damage to epithelial-endothelial integrity, highlighting the need for complementary strategies targeting barrier repair and tissue regeneration. Importantly, the ability to resolve such distinct functional outcomes demonstrates the utility of the system for mechanistic studies of therapeutic interventions at the alveolar interface.

From a broader perspective, this work highlights the importance of interfacial material design in organ-on-a-chip systems. By linking membrane composition to degradation kinetics, structural evolution, and transport behavior, the platform provides a framework for engineering dynamic biological interfaces that more closely replicate native tissue environments. This concept extends beyond lung models and may be applicable to other barrier tissues where structure–function relationships evolve over time.

Future studies can further expand the capabilities of this system. Incorporation of additional cell types, such as fibroblasts or immune cells, would enable investigation of matrix remodeling, immune responses, and chronic injury processes. Integration of real-time sensing modalities, including barrier resistance and oxygen transport, could provide continuous functional readouts. In addition, extending the culture duration and incorporating repeated or low-dose exposures may allow modeling of chronic particulate exposure and disease progression. Finally, systematic variation of membrane composition and degradation kinetics could provide further insight into how dynamic interfacial properties regulate cellular responses.

In conclusion, we present a biodegradable PLGA/PCL membrane-based lung alveoli-on-a-chip that combines dynamic structural evolution, physiologically relevant transport, and mechanical stimulation to model particulate-induced alveolar injury. By enabling layer-resolved analysis of toxicity and selective therapeutic modulation, this platform provides a versatile and translatable tool for studying air pollution-related lung disease and for evaluating candidate interventions.

## 4. Material and Methods

### 4.1 PLGA/PCL membrane fabrication

All chemicals were purchased from Sigma-Aldrich (PLGA: P219, PCL: 440744, camphene: 456055) and used as received. A 10 wt% PLGA/PCL solution was prepared by dissolving the polymers in acetone at 60 °C under magnetic stirring for 3 hours until fully homogenized. Camphene, used as a porogen, was then. added at 100 wt% relative to the polymer content and mixed at 45 °C for an additional 1 hour. The resulting polymer-camphene solution was cooled to room temperature prior to membrane fabrication.

Polymer membranes were produced by spin-coating the polymer-camphene solution onto a silicon wafer at 400 rpm for 5 seconds. The coated wafers were immediately immersed in distilled water at room temperature for 10 minutes to induce nonsolvent-driven phase separation and solidification of the polymer matrix, forming a transient pore template. Subsequently, camphene was selectively removed by freeze-drying for 3 hours, yielding an interconnected porous PLGA/PCL membrane.

### 4.2 Membrane characterization

The surface morphology and internal porous structure of the PLGA/PCL membranes were characterized using scanning electron microscopy (SEM; Quanta 600 FEG, FEI Company, USA). For cross-sectional observation, the membranes were cryo-fractured in liquid nitrogen prior to imaging. Samples were fixed in 2.5% glutaraldehyde for 1 hour at room temperature, rinsed three times with phosphate-buffered saline (PBS), and post-fixed in 1% osmium tetroxide (OsO₄) for 1 hour at 4 °C. The specimens were then dehydrated through a graded ethanol series (50–100%) and treated with hexamethyldisilazane (HMDS, Sigma-Aldrich) for 15 minutes, followed by air drying. Surface porosity was quantified from SEM images using ImageJ software. For each condition, four independent SEM micrographs were analyzed, and porosity was calculated based on threshold-based area measurements.

### 4.3 Assessment of membrane degradation behavior

To assess membrane degradation, PLGA/PCL membranes were incubated in PBS at 37 °C for predetermined time intervals. After incubation, the samples were rinsed with distilled water and freeze-dried for 3 hours to remove residual moisture within the pores. The dry weight of the degraded membrane (Wₜ) was then recorded. The degradation ratio was calculated according to the following equation: 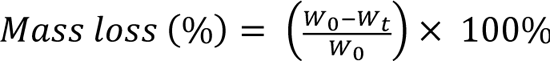, where W_0_ represents the initial dry weight of the membrane.

### 4.4 Tensile strength measurement

The mechanical properties of the PLGA/PCL membranes were evaluated by uniaxial tensile testing. Specimens were elongated at a constant crosshead speed of 10 mm/min using a universal testing machine (MTS Criterion® 42) fitted with a 50 N load cell. Stress-strain data were continuously acquired, and the apparent Young’s modulus was calculated from the slope of the initial linear portion of the stress-strain curves. Membrane thickness was measured from cross-sectional SEM images to accurately normalize the applied force and determine stress values.

### 4.5 Fourier transform infrared spectroscopy

Fourier transform infrared (FT-IR) spectroscopy (PerkinElmer Inc., USA) was used to characterize the chemical structure of the PLGA/PCL membranes. Spectra of PLGA/PCL membranes were acquired over the range of 600–2000 cm⁻¹, with 32 scans collected per sample to obtain adequate spectral resolution and signal quality.

### 4.6 Permeability Measurement

Membrane permeability was assessed by first removing the culture medium and gently washing the device with PBS. Next, 1 mL of phenol red–free culture medium was added to the basolateral chamber, while 0.5 mL of FITC-dextran solution (4 kDa; Sigma-Aldrich) was introduced into the apical chamber. The chip was then incubated at 37 °C, and samples were collected from the basolateral chamber at 0, 10, 20, 30, 40 minutes into a black 96-well plate. Fluorescence intensity was quantified using a microplate reader (SpectraMax i3x) with excitation and emission wavelengths set to 490 nm and 525 nm, respectively. The apparent permeability coefficient (P_app_, cm s⁻¹) was calculated according to the equation:

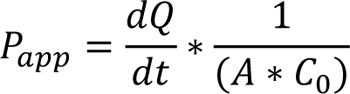

where dQ/dt is the rate of FITC-dextran transport across the membrane, A denotes the effective membrane area (cm²), and C₀ represents the initial concentration of the fluorescent tracer in the apical chamber (mg mL⁻¹).

### 4.7 Device assembly

To assemble the lung alveoli-on-a-chip device, PLGA/PCL membrane was initially bonded to the apical PDMS chamber (surface area: 1.13 cm²) using uncured PDMS as an adhesive and cured overnight at 40 °C to achieve robust attachment. The apical chamber with the integrated membrane was subsequently aligned and bonded to the basolateral PDMS chamber (area: 2.01 cm²) using a thin layer of liquid PDMS, followed by curing at 40 °C overnight to ensure complete and stable sealing.

### 4.8 Cell culture

For cell culture, the devices were sterilized by UV irradiation for 3 hours following spraying with 70% ethanol. Prior to cell seeding, the membranes were coated with a 0.1% gelatin solution (Cell Biologics) at 37 °C for 5 minutes to enhance cell adhesion. For endothelial layer formation, the devices were inverted and human lung microvascular endothelial cells (HMVECs, Lonza) were seeded at a density of 2 × 10⁵ cells cm⁻². After 4 hours of incubation to allow cell attachment, the devices were returned to the upright orientation, and human primary alveolar epithelial cells (HPAECs, Cell Biologics) were introduced into the apical chamber at a density of 5 × 10⁵ cells cm⁻².

The HPAEC-HMVEC co-cultures were maintained in endothelial growth medium (EGM-2, Lonza, CC-3202) under liquid–liquid culture conditions at 37 °C and 5% CO₂ for up to 7 days to facilitate barrier formation. On day 7, the apical medium was removed to establish ALI condition. After 24 hours to allow ALI stabilization, cyclic mechanical stimulation was applied to recapitulate physiological breathing-mimetic motions.

A flexible PDMS lid, prepared by spin coating, was attached to the top of the apical chamber. Periodic compression of the PDMS lid was achieved using a solenoid actuator (Adafruit Mini Push-pull solenoid) driven by an Arduino microcontroller. The actuator operated at a frequency of 0.33 Hz, inducing cyclic deformation of the membrane.

Assuming that the deformed membrane adopts an approximately partial spherical geometry, the linear strain (*ΔL/L*) was estimated using the following equation, where *a* denotes the membrane radius and *h* represents the measured axial deflection during compression.

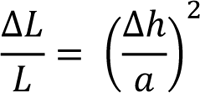

The axial deflection during compression was calculated by measuring the volume displacement of the fluid in the basolateral chamber. During cell culture, the medium in the basolateral chamber was replaced every 2 days.

### 4.9 Live/Dead cell viability assay

Cell viability was evaluated using a Live/Dead Cell Viability and Cytotoxicity Assay Kit (Biotium, USA). After gentle washing with PBS, the samples were incubated at 37 °C for 30 minutes in a staining solution consisting of 2 µM calcein AM and 4 µM ethidium homodimer III (EthD-III) diluted in culture medium. The samples were then rinsed twice with PBS and placed in fresh medium prior to imaging. Fluorescence images were obtained using a Leica DMi8 microscope with a 20× objective, and cell viability was quantified by analyzing four randomly selected fields of view for each sample.

### 4.10 Immunofluorescence staining

Cells grown on the membranes were rinsed three times with PBS and fixed in 4% paraformaldehyde (Sigma-Aldrich) for 15 minutes at room temperature. The membranes were then sectioned, and the fixed cells were permeabilized with 0.1% Triton X-100 in PBS for 10 minutes at room temperature. The samples were subsequently blocked with PBS containing 5% donkey serum for 30 minutes at room temperature. Primary antibodies against ZO-1 (Rockland, 600-401-GU7, 1:400), VE-cadherin (Cell Signaling Technology, 2500, 1:200), or phospho-histone H2A.X (Cell Signaling Technology, 1:200) were prepared in blocking buffer and incubated with the samples overnight at 4 °C. After thorough washing with PBS, the samples were incubated with the corresponding secondary antibodies for 1 hour at room temperature. Nuclei were counterstained with DAPI after secondary antibody labeling. Fluorescence images were captured using a Leica DMi8 fluorescence microscope.

### 4.11 Particle size analysis of DPM

The particle size distribution of DPM was characterized using a combination of dynamic light scattering (DLS) and laser diffraction techniques to capture nano to micro-scale particle populations. Measurements were performed using a Zetasizer Nano ZS and a Malvern Panalytical Mastersizer, respectively, at the Materials Characterization Laboratory at Pennsylvania State University.

For nano-scale particle characterization, the Zetasizer Nano ZS was employed, which operates based on the principle of DLS and has a typical measurement range of approximately 0.3 nm to 10 μm. Prior to measurement, a representative DPM sample was dispersed in isopropanol as the dispersing medium. The suspension was ultrasonicated for approximately 5 minutes to promote deagglomeration and achieve preliminary dispersion. A small aliquot of the dispersed sample was then transferred into a 12 mm outer-diameter square polystyrene cuvette suitable for organic solvents and allowed to remain undisturbed for 2 minutes to stabilize the dispersion prior to analysis. In DLS measurements, thermally induced collisions between solvent molecules and suspended DPM particles cause the particles to undergo Brownian motion. When illuminated by a laser, these randomly moving particles scatter light, producing time-dependent intensity fluctuations. Analysis of these fluctuations yields the translational diffusion coefficient of the particles, which is converted to particle size using the Stokes-Einstein relationship. The diameter reported by DLS corresponds to the hydrodynamic diameter, representing the effective size of particles as they diffuse within the liquid medium. Data acquisition and analysis were conducted using the Zetasizer software (version 8.01).

Particle size distributions were also measured using the Mastersizer, which employs laser diffraction and covers a measurement range of approximately 0.01–3500 μm. Laser diffraction is particularly effective for larger particles, as they scatter light at higher intensities and lower angles compared to nano-scale particles. The complementary use of the Zetasizer and Mastersizer allowed for comprehensive characterization of particle size distribution across multiple orders of magnitude.

### 4.12 Hydrogen peroxide and cytokine quantification

The concentrations of H₂O₂ and multiple cytokines were quantified using a Hydrogen Peroxide Assay Kit (Sigma-Aldrich) and a Multiplex Human Cytokine ELISA Kit (Anogen), respectively, in accordance with the manufacturers’ instructions.

### 4.13 Quantification of γ-H2AX-positive cells

The percentage of γ-H2AX-positive cells was calculated by normalizing the number of γ-H2AX-stained cells to the total number of cells within a defined region of interest. For each sample, four independent fluorescence images were collected, and their values were averaged to obtain the mean γ-H2AX positivity.

### 4.14 Statistics

Statistical analyses were performed using Student’s t-test for comparisons between two groups. For experiments with three or more groups, one-way or two-way analysis of variance (ANOVA) was employed, followed by Tukey’s post hoc test for multiple comparisons. Differences were considered statistically significant at p < 0.05.

---

**Figure S1.**
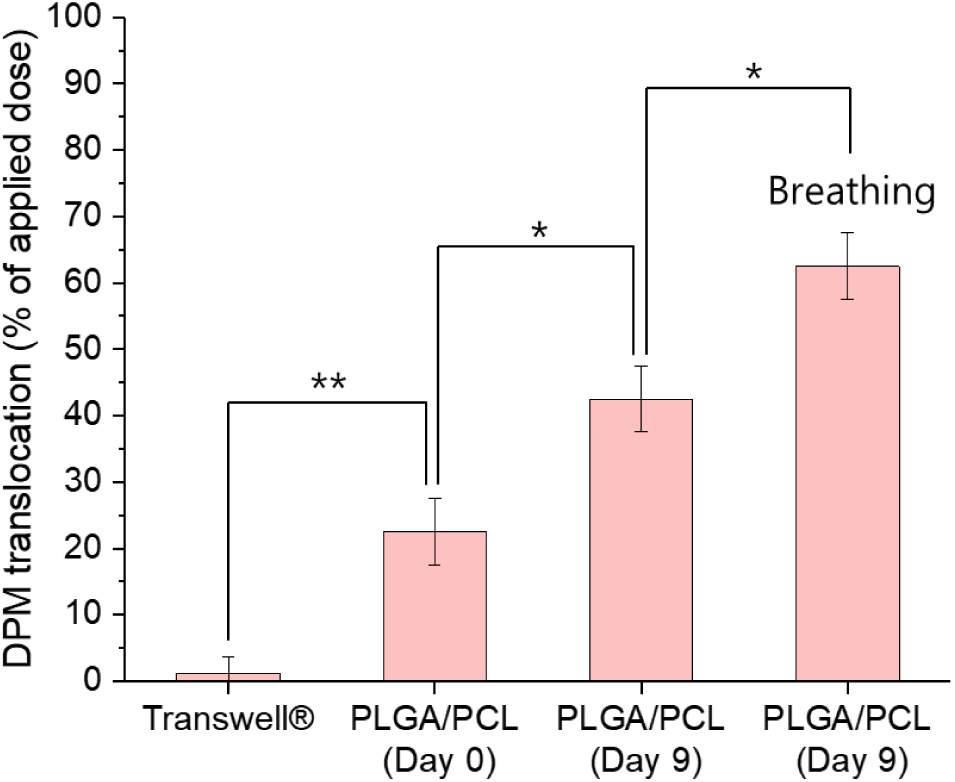
Percentage of DPM translocated across the membrane 24 hours after apical administration of 100 µg DPM (n = 4).

## Notes

### Competing Interest Statement

The authors have declared no competing interest.

## References

[1] 2025 Global Burden of Disease 2023.

[2] 2025 Burden of 375 diseases and injuries, risk-attributable burden of 88 risk factors, and healthy life expectancy in 204 countries and territories, including 660 subnational locations, 1990-2023: a systematic analysis for the Global Burden of Disease Study 2023 Lancet 406 1873–922

[3] Azam S, Liu S, Bhattacharyya S and Zheng S 2024 Assessing the hazard of diesel particulate matter (DPM) in the mining industry: A review of the current state of knowledge International Journal of Coal Science & Technology 11

[4] Reis H, Reis C, Sharip A, Reis W, Zhao Y, Sinclair R and Beeson L 2018 Diesel exhaust exposure, its multi-system effects, and the effect of new technology diesel exhaust Environ Int 114 252–65

[5] Steiner S, Bisig C, Petri-Fink A and Rothen-Rutishauser B 2016 Diesel exhaust: current knowledge of adverse effects and underlying cellular mechanisms Arch Toxicol 90 1541–53

[6] Miller MR, McLean SG, Duffin R, Lawal AO, Araujo JA, Shaw CA, Mills NL, Donaldson K, Newb DE and Hadoke PW 2013 Diesel exhaust particulate increases the size and complexity of lesions in atherosclerotic mice Particle and Fibre Toxicology 10 61

7. [7] Mills NL, Tornqvist H, Robinson SD, Gonzalez M, Darnley K, MacNee W, Boon NA, Donaldson K, Blomberg A, Sandstrom T and Newby DE 2005 Diesel exhaust inhalation causes vascular dysfunction and impaired endogenous fibrinolysis Circulation 112 3930–6

[8] Pope CA, 3rd, Bhatnagar A, McCracken JP, Abplanalp W, Conklin DJ and O’Toole T 2016 Exposure to Fine Particulate Air Pollution Is Associated With Endothelial Injury and Systemic Inflammation Circ Res 119 1204–14

[9] Ristovski ZD, Miljevic B, Surawski NC, Morawska L, Fong KM, Goh F and Yang IA 2012 Respiratory health effects of diesel particulate matter Respirology 17 201–12

[10] Sydbom A, Blomberg A, Parnia S, Stenfors N, Sandström T and Dahlen SE 2001 Health effects of diesel exhaust emissions European Respiratory Journal 17 733–46

[11] McCreanor J, Cullinan P, Nieuwenhuijsen MJ, Stewart-Evans J, Malliarou E, Jarup L, Harrington R, Svartengren M, Han I-K, Ohman-Strickland P, Chung KF and Zhang J 2007 Respiratory effects of exposure to diesel traffic in persons with asthma New England Journal of Medicine 357 2348–58

[12] Duval K, Grover H, Han LH, Mou Y, Pegoraro AF, Fredberg J and Chen Z 2017 Modeling Physiological Events in 2D vs. 3D Cell Culture Physiology (Bethesda) 32 266–77

[13] Upadhyay S and Palmberg L 2018 Air-Liquid Interface: Relevant In Vitro Models for Investigating Air Pollutant-Induced Pulmonary Toxicity Toxicol Sci 164 21–30

[14] Ingber DE 2020 Is it Time for Reviewer 3 to Request Human Organ Chip Experiments Instead of Animal Validation Studies? Adv Sci (Weinh) 7 2002030

[15] Bhatia SN and Ingber DE 2014 Microfluidic organs-on-chips Nat Biotechnol 32 760–72

[16] Zhang B, Korolj A, Lai BFL and Radisic M 2018 Advances in organ-on-a-chip engineering Nature Reviews Materials 3 257–78

[17] Ringquist R, Bhatia E, Chatterjee P, Maniar D, Fang Z, Franz P, Kramer L, Ghoshal D, Sonthi N, Downey E, Canlas J, Ochal A, Agarwal S, Cuellar V, Harrigan G, Coskun AF, Singh A and Roy K 2025 An immune-competent lung-on-a-chip for modelling the human severe influenza infection response Nat Biomed Eng 1–23

[18] Yadav S, Fujimoto K, Takenaga T, Takahashi S, Muramoto Y, Mikawa R, Noda T, Gotoh S and Yokokawa R 2025 Isogenic induced-pluripotent-stem-cell-derived airway- and alveolus-on-chip models reveal specific innate immune responses Nat Biomed Eng 1–15

[19] Dasgupta Q, Jiang A, Wen AM, Mannix RJ, Man Y, Hall S, Javorsky E and Ingber DE 2023 A human lung alveolus-on-a-chip model of acute radiation-induced lung injury Nat Commun 14 6506

[20] Huang D, Liu T, Liao J, Maharjan S, Xie X, Perez M, Anaya I, Wang S, Tirado Mayer A, Kang Z, Kong W, Mainardi VL, Garciamendez-Mijares CE, Garcia Martinez G, Moretti M, Zhang W, Gu Z, Ghaemmaghami AM and Zhang YS 2021 Reversed-engineered human alveolar lung-on-a-chip model Proc Natl Acad Sci U S A 118 e2016146118

[21] Doryab A, Taskin MB, Stahlhut P, Schröppel A, Wagner DE, Groll J and Schmid O 2020 A Biomimetic, Copolymeric Membrane for Cell-Stretch Experiments with Pulmonary Epithelial Cells at the Air-Liquid Interface Advanced Functional Materials 31

[22] Baptista D, Teixeira LM, Birgani ZT, van Riet S, Pasman T, Poot A, Stamatialis D, Rottier RJ, Hiemstra P S, Habibovic P, van Blitterswijk C, Giselbrecht S and Truckenmuller R 2021 3D alveolar in vitro model based on epithelialized biomimetically curved culture membranes Biomaterials 266 120436

[23] Garnock-Jones KP 2015 Roflumilast: A Review in COPD Drugs 75 1645–56

[24] Giembycz MA and Field SK 2010 Roflumilast: first phosphodiesterase 4 inhibitor approved for treatment of COPD Drug Des Devel Ther 4 147–58

[25] Li H, Zuo J and Tang W 2018 Phosphodiesterase-4 Inhibitors for the Treatment of Inflammatory DiseasesFront Pharmacol 9 1048

[26] Knudsen L and Ochs M 2018 The micromechanics of lung alveoli: structure and function of surfactant and tissue components Histochem Cell Biol 150 661–76

[27] Nair LS and Laurencin CT 2007 Biodegradable polymers as biomaterials Progress in Polymer Science 32 762–98

[28] Malikmammadov E, Tanir TE, Kiziltay A, Hasirci V and Hasirci N 2018 PCL and PCL-based materials in biomedical applications Journal of Biomaterials Science, Polymer Edition 29 863–93

[29] Guillen GR, Pan Y, Li M and Hoek EMV 2011 Preparation and Characterization of Membranes Formed by Nonsolvent Induced Phase Separation: A Review Industrial & Engineering Chemistry Research 50 3798–817

[30] Loffler RJG, Hanczyc MM and Gorecki J 2022 A camphene-camphor-polymer composite material for the production of superhydrophobic absorbent microporous foams Sci Rep 12 243

[31] Hough RF, Bhattacharya S and Bhattacharya J 2018 Crosstalk signaling between alveoli and capillaries Pulm Circ 8 2045894018783735

[32] Wagner PD 2015 The physiological basis of pulmonary gas exchange: implications for clinical interpretation of arterial blood gases Eur Respir J 45 227–43

[33] Rowan SC, Rochfort KD, Piouceau L, Cummins PM, O’Rourke M and McLoughlin P 2018 Pulmonary endothelial permeability and tissue fluid balance depend on the viscosity of the perfusion solution Am J Physiol Lung Cell Mol Physiol 315 L476–L84

[34] Brune K, Frank J, Schwingshackl A, Finigan J and Sidhaye VK 2015 Pulmonary epithelial barrier function: some new players and mechanisms Am J Physiol Lung Cell Mol Physiol 308 L731–45

[35] Salvator H, Buenestado A, Brollo M, Naline E, Victoni T, Longchamp E, Tenor H, Grassin-Delyle S and Devillier P 2020 Clinical Relevance of the Anti-inflammatory Effects of Roflumilast on Human Bronchus: Potentiation by a Long-Acting Beta-2-Agonist Front Pharmacol 11 598702

[36] Milara J, Armengot M, Banuls P, Tenor H, Beume R, Artigues E and Cortijo J 2012 Roflumilast N-oxide, a PDE4 inhibitor, improves cilia motility and ciliated human bronchial epithelial cells compromised by cigarette smoke in vitro Br J Pharmacol 166 2243–62

[37] Mata M, Martinez I, Melero JA, Tenor H and Cortijo J 2013 Roflumilast inhibits respiratory syncytial virus infection in human differentiated bronchial epithelial cells PLoS One 8 e69670

[38] Hohne K, Schliessmann S J, Kirschbaum A, Plones T, Muller-Quernheim J, Tenor H and Zissel G 2012 Roflumilast-N-oxide induces surfactant protein expression in human alveolar epithelial cells type II PLoS One 7 e38369

[39] Buenestado A, Grassin-Delyle S, Guitard F, Naline E, Faisy C, Israel-Biet D, Sage E, Bellamy J F, Tenor H and Devillier P 2012 Roflumilast inhibits the release of chemokines and TNF-alpha from human lung macrophages stimulated with lipopolysaccharide Br J Pharmacol 165 1877–90

